# Single-cell analyses reveal scyphozoan conserved and specific genes contribution during tissue regeneration in two jellyfish

**DOI:** 10.1101/2025.04.29.650939

**Authors:** Yiqian Li, Sean T.S. Law, Wenyan Nong, Wai Lok So, Yichun Xie, Thomas C.N. Leung, Tse Ho Li, Joyce Tse, Ho Yin Yip, Oli Jin, Jordan Zhang, Apple PY Chui, Kwok Fai Lau, Akbar John, Zhen-peng Kai, William G. Bendena, Alexander Hayward, Yingying Wei, Ting Fung Chan, Sai Ming Ngai, Jerome HL Hui

**Affiliations:** School of Life Sciences, Simon F.S. Li Marine Science Laboratory, State Key Laboratory of Agrobiotechnology, Institute of Environment, Energy and Sustainability, The Chinese University of Hong Kong, Hong Kong SAR, China; School of Life Sciences, State Key Laboratory of Agrobiotechnology, The Chinese University of Hong Kong, Hong Kong SAR, China; Dovetail Genomics, USA; School of Life Sciences, Simon F.S. Li Marine Science Laboratory, The Chinese University of Hong Kong, Hong Kong SAR, China; School of Life Sciences, The Chinese University of Hong Kong, Hong Kong SAR, China; College of Marine Science and Aquatic Biology, University of Khorfakkan, Sharjah, United Arab Emirates; Shanghai Institute of Technology, Shanghai, China; Department of Biology, Queen’s University, Canada; University of Exeter, United Kingdom; Department of Statistics, The Chinese University of Hong Kong, Hong Kong SAR, China

**Author notes:** correspondence =. contributed equally (Yiqian Li, Sean T.S. Law, Wenyan Nong).

## Abstract

The phylum Cnidaria is the outgroup of Bilateria and includes sea anemones, corals, hydroids, and jellyfish. Cnidarians play crucial ecological roles in marine ecosystems, including the formation of highly diverse and productive coral reefs, and acting as important predator and prey species. Cnidarians are also well known for their remarkable regeneration capacities. Here, we report single-cell RNA sequencing during bell regeneration in two species of scyphozoans or true jellyfish, *Aurelia coerulea* and *Rhopilema esculentum*. We delineated 12 cell populations in *Aurelia* and *Rhopilema*, and revealed their respective marker genes and enriched gene pathways. Conserved transcription factor *Otx* and Wnt/ β-catenin signalling pathway genes were identified during regeneration processes. Additionally, we discovered two conserved, sequentially activated modules during regeneration, with stem cells, gastrodermal cells, neural cells, and secretory gland cells modulated in the first phase, followed by cnidocytes in the second phase. Further comparison among cnidarian genomes identified a suite of scyphozoan lineage-specific genes, a subset of which were frequently significantly expressed in cnidocytes in both jellyfish species during the first phase of regeneration. This study reveals key insights into the evolution and contribution of conserved and novel genes to the formation of lineage-specific genetic networks and biological processes.

## Introduction

Animals display a fascinating diversity of body plans and cellular mechanisms that facilitate adaptation to the environment. The bilaterians comprise most extant animal species, and to understand how bilaterians evolved, it is essential to consider their immediate outgroup – the phylum Cnidaria. The Cnidaria contains ∼11,000 described species including sea anemones, corals, hydroids, and jellyfish, that play important ecological roles in marine, and less commonly, freshwater environments. Macroevolutionary trends in gene repertoire evolution have been analysed across many cnidarians over the last decade, with representative genomes of key cnidarian groups assembled, including anthozoans or ‘sea anemones and corals’ (Putnam et al 2007; Shinzato et al 2011; Baumgarten et al 2015; Shum et al 2022; Yu et al 2022), hydrozoans or ‘hydroids’ (Chapman et al 2010; Leclere et al 2019), myxozoans (Chang et al 2015), and scyphozoans or ‘true jellyfish’ (Gold et al 2019; Khalturin et al 2019; Nong et al 2020) and cubozoans or ‘box jellyfish’ (Liegertová et al 2015; Khalturin et al 2019; Ohdera et al 2019). These data have provided important insights into the evolutionary pathways of both bilaterians and cnidarians.

Regeneration refers to the ability to restore a lost body part, and represents an important process occurring in many animals. It likely contributes to evolutionary success, given the importance associated with replacing lost body parts (Bideau et al 2021; Gurtner et al 2008). Nevertheless, regenerative capacities vary across lineages in both cnidarians and bilaterians. For instance, while bilaterians such as amphibians, earthworms, reptiles, and zebrafish can heal wounds and regenerate certain lost organs (Liu et al 2015; Nowoshilow et al 2018; Shao et al 2020), planarian flatworms can regenerate almost the whole body, similarly to some cnidarians (i.e. hydrozoans and anthozoans; Vila-Farre et al 2023; Vogg et al 2019; DuBuc et al 2014; Sinigaglia et al 2020; Layden et al 2016). Yet, compared to many other cnidarians lineages, such as anthozoans and hydrozoans, the study of regeneration in scyphozoans has been relatively neglected.

Scyphozoans play important ecological roles throughout the oceans in different parts of the world and are well-known to humans due to their often-painful stings, the occurrence of largescale jellyfish blooms, and their use in aquaculture as food in some countries. In contrast to the colonial hydrozoans, scyphozoans have both asexually sessile polyps and dispersive and sexually reproductive medusae in their life cycles. Regenerative capacities have been demonstrated in certain species of scyphozoans at different life stages (Zeleny 1907; Steinberg 1963; Curtis and Cowden 1974; Stierwald et al 2004; Gamero-Mora et al 2019; Abrams et al 2015; Abrams et al 2021). However, the underlying cellular and genetic components contributing to scyphozoan regeneration remain poorly elucidated.

Here, we present the first single-cell transcriptomics analysis of regeneration in Scyphozoa, focussing on two species, the moon jellyfish *Aurelia coerulea* (Semaeostomeae) and the edible jellyfish *Rhopilema esculentum* (Rhizostomeae). Our findings provide insights into the evolutionary pathways underlying regeneration in scyphozoans.

## Results

### Resequencing of a new moon jellyfish genome

Genomic resources for moon jellyfish *Aurelia aurelia* and edible jellyfish *Rhopilema esculentum* were generated previously (Figure 1A-B) (Gold et al 2019; Nong et al 2020). However, genomes assembled from different populations of the cosmopolitan moon jellyfish show considerable variation (Khalturin et al. 2019; Dong et al. 2024). Correspondingly, in sequence alignments between single-cell sequencing reads generated here and published genomes, revealed sequence similarity of 96-98% for *Rhopilema*, but 39%-95% for *Aurelia* (Supplementary Table 2,6). The *Aurelia* draft genome originates from a specimen in California (Gold et al 2019), and features an unusually high number of BUSCO duplicates and relatively short scaffold sizes (given today’s standards). Thus, given these considerations, and the low rate of mapping observed for our *Aurelia* reads, we decided to generate a new assembly for a male moon jellyfish, which was obtained from Hong Kong (Supplementary Figure 1). The new genome, *Aurelia* (CUHK_M) features an assembly size of 449.7Mb (Figure 1B), and is comparable to the published *Aurelia* genome size and duplicated BUSCO numbers, but it features considerably improved sequence continuity (contig N50 = 8.9Mb; scaffold N50 = 22.4Mb; Figure 1C; Supplementary Figure 1, Supplementary Table 1-3). We also achieved mapping rates of 90-92% for single-cell sequencing reads, considerably improving on efforts based on the other *Aurelia* draft genome (Supplementary Table 1-4).

**Figure 1.**
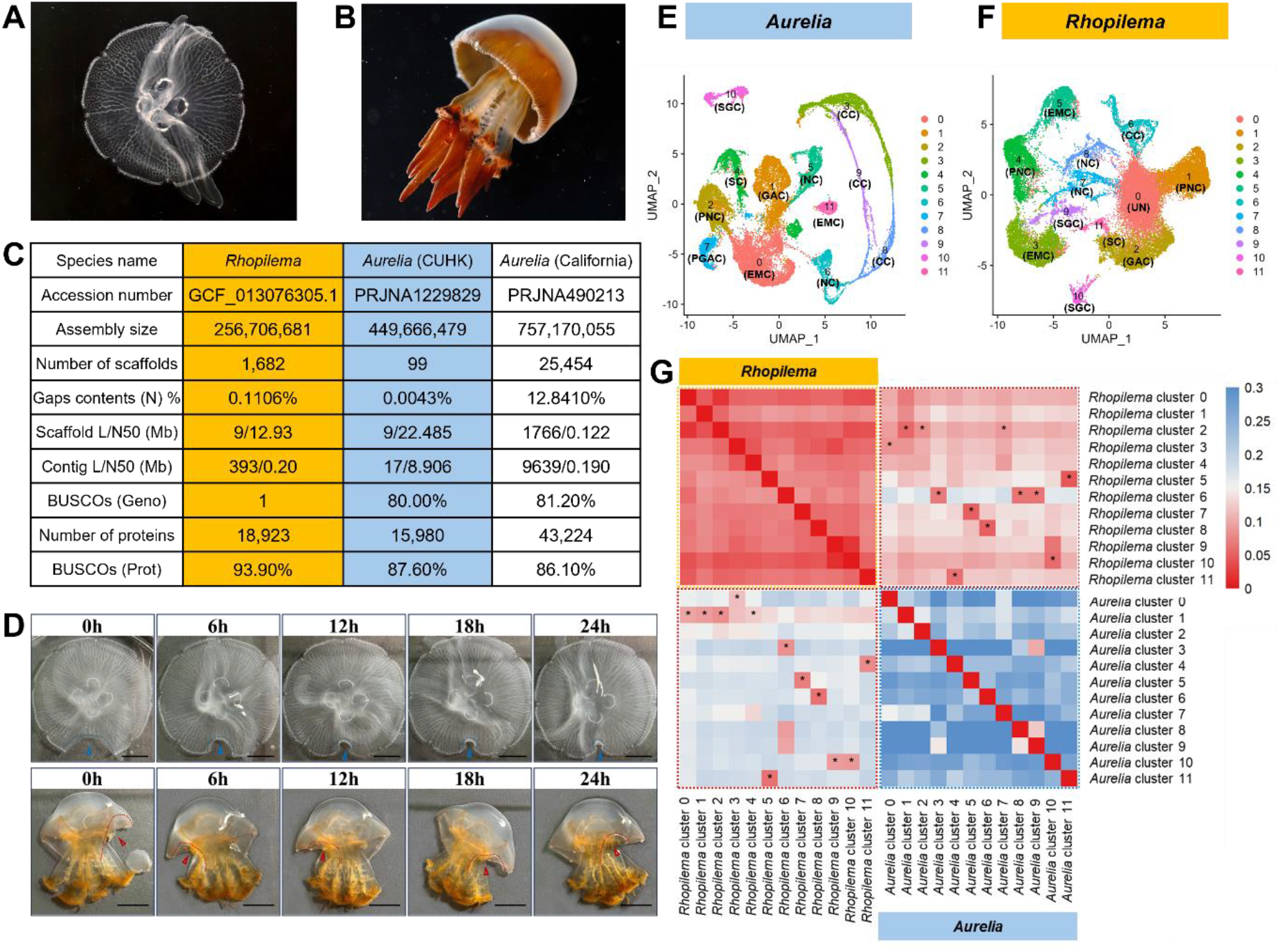
**A)** Moon jellyfish *Aurelia*. **B)** Edible jellyfish *Rhopilema*. **C)** Genome assembly quality. **D)** Bell regeneration of *Aurelia* (upper) and *Rhopilema* (lower). The arrowheads depit the amputated regions. Scale bar = 1cm. **E-F)** UMAP visualization of twelve cell clusters of *Aurelia* (E) and *Rhopilema* (F). **G)** Marker genes comparison between *Aurelia* and *Rhopilema*.

### *Aurelia* and *Rhopilema* cell type atlas

Single-cell suspensions were generated by regenerating bell tissues of *Aurelia* and *Rhopilema* at five timepoints (0h, 6h, 12h, 18h, 24h) for single-cell RNA sequencing (scRNA-seq; Figure 1D; Supplementary Table 5,8; Supplementary Figure 2). The resultant scRNA-seq libraries contained an average of 53,280 reads/cell and 745 genes/cell for *Aurelia*, and an average of 35,597 reads/cell and 709 genes/cell for *Rhopilema*, which is comparable to other published cnidarian scRNA-seq studies (Chari et al., 2021; Hu et al., 2020; Levy et al., 2021; Sebé-Pedrós et al., 2018; Siebert et al., 2019, Dong et al., 2024). Using uniform manifold approximation and projection (UMAP) analyses on ∼27,000 and ∼31,000 single-cell profiles, 12 cell clusters with unique gene expression profiles were identified in both *Aurelia* and *Rhopilema* (Figure 1E-F).

**Figure 2.**
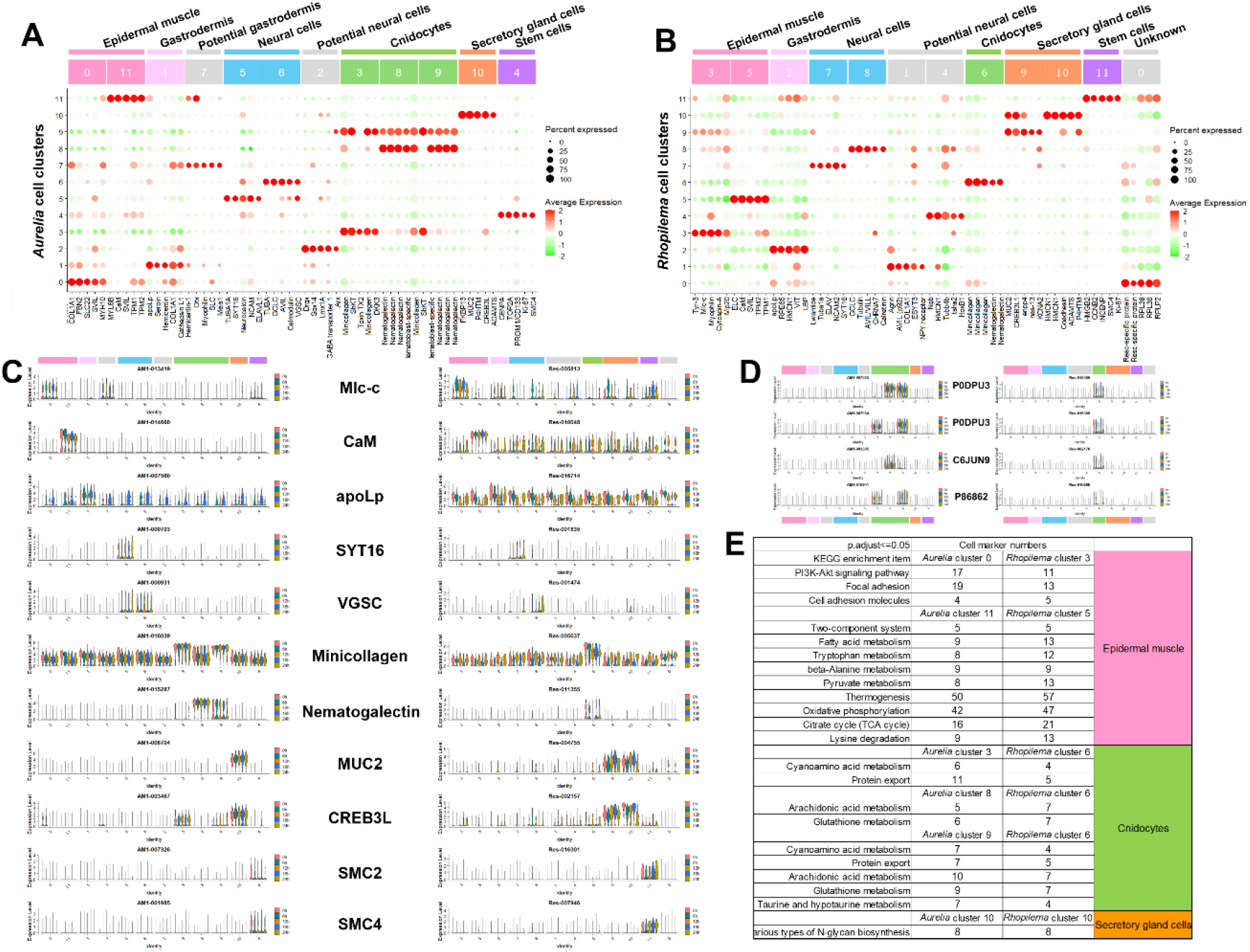
**A-B)** Marker genes of *Aurelia* cell clusters (A) and *Rhopilema esculentum* cell clusters (B). Pink: Epidermal muscle (EM) and Gastrodermis (GAS); Blue: Neural cells (NC) and Potential neural cells (PNC); Green: Cnidocytes (CC); Orange: Secretory.gland cells (SGC); and Purple: Stem/germline cells (SC). **C)** Expression of cell-type-specific marker genes in different cell types of *Aurelia* (left) and *Rhopilema* (right). **D)** Expression of conserved cnidocyte toxin genes in *Aurelia* (left) and *Rhopilema* (right). **E)** Enriched KEGG pathways of genes in the same cell types between *Aurelia* and *Rhopilema*.

To generate the cell atlas, corresponding highly expressed marker genes were first identified and compared to known cnidarian cell types, including hydrozoans *Clytia hemisphaerica* and *Hydra vulgaris*; sea anemone *Nematostella vectensis*, and corals *Xenia sp*. and *Stylophora pistillata* (Chari et al., 2021; Hu et al., 2020; Levy et al., 2021; Siebert et al., 2019)(Supplementary Table 9-10; Supplementary Figure 6-11). In sum, a total of 8 cell types including cnidocytes/nematocytes (CC; *Aurelia* clusters 3,8,9, *Rhopilema* cluster 6), epidermal muscle (EM; *Aurelia* cluster 0,11, *Rhopilema* cluster 3,5), gastrodermis (GDC; *Aurelia* cluster 1, *Rhopilema* cluster 2), potential gastrodermis (PGDC), neural cells (NC; *Aurelia* clusters 5,6, *Rhopilema* clusters 7,8), potential neural cells (PNC), secretory gland cells (SGC; *Aurelia* cluster 10, *Rhopilema* clusters 9,10), and stem/germline cells (SC; *Aurelia* cluster 4, *Rhopilema* cluster 11) could be assigned to all gene clusters in both *Aurelia* and *Rhopilema* (Figure1E-G)(Supplementary Figure 4-5). In addition, we also performed mass spectrometry for cnidocysts, revealing conserved expression of 4 potential toxins in the assigned cnidocytes/nematocytes of both *Aurelia* and *Rhopilema* (Figure 2D; Supplementary Figure 12). Thus, a combination of signature marker gene expression and mass spectrometry allowed us to solidly assign cell identities (Figure 2A-C, Supplementary Figure 4-5).

Sesquiterpenoid hormone and its biosynthetic pathway genes were previously described in jellyfish (Nong et al 2020). Utilising the cell atlas generated here, we found that genes involved in the sesquiterpenoid hormone biosynthetic pathway, including *AACT, HMGCS, HMGCR, MVK, MDC, FNTB, FNTA, ZMPSTE24*, and *IMCT* are expressed at different levels among cell types (Supplementary Figure 13). Among these genes, *AACT* and *ICMT* have the highest expression level in most cell types in both species.

We further tested if any functional gene ontology terms, including those from the GO, KEGG and KOG databases were enriched in these different cell types (Figure 2E, Supplementary Table 11-12; Supplementary Figure 14). As shown in Figure 2E, conserved gene pathways were also identified in most assigned cell types, including cnidocysts/nematocysts, epidermal muscle, secretory gland cells, neural cells and stem cells, between *Aurelia* and *Rhopilema*.

### Gene modulation during tissue regeneration in *Aurelia* and *Rhopilema*

To assess the interplay of cellular and underlying molecular processes contributing to regeneration in *Aurelia* and *Rhopilema*, we compared scRNA-seq data generated at five distinct time points. Pseudotemporal analysis revealed similar but divergent reprogramming of cell types during regeneration (Figure 3A; Supplementary Figure 15). For instance, in both species, modulation of stem cells occurred first, followed by modulation of gastrodermal cells, secretory gland cells and neural cells in the early phase of regeneration, while cnidocytes modulated in the second phase of regeneration. Epidermal muscle cells differentiated concurrently with gastrodermis, neural cells, and secretory gland cells in early phase of regeneration in *Aurelia*, and yet, these cell types developed in the later phase of regeneration in *Rhopilema*.

**Figure 3.**
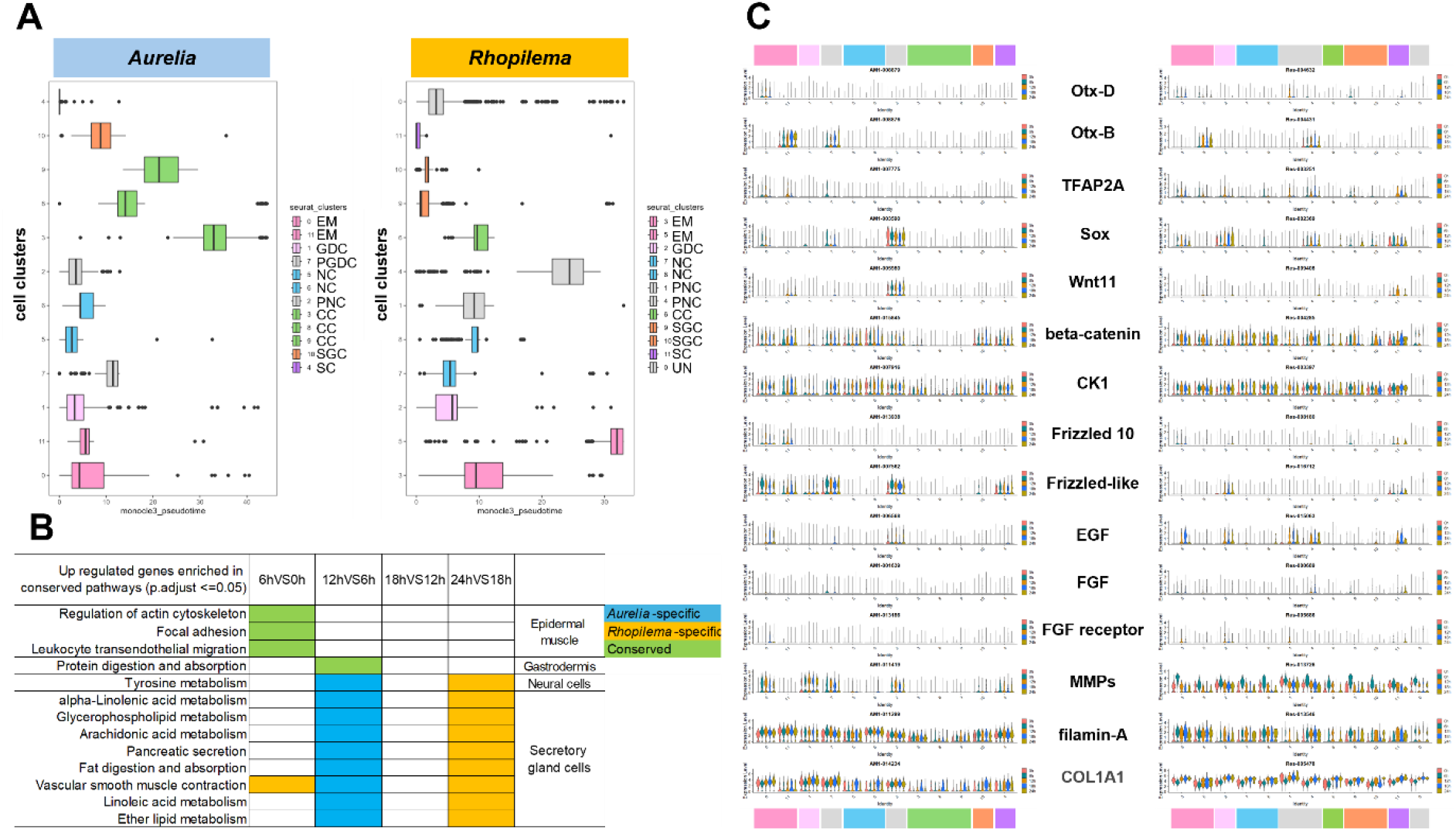
**A)** Pseudotemporal cell ordering of bell regeneration in *Aurelia* (left) and *Rhopilema* (right). **B)** Enriched KEGG pathways of conserved up-regulated genes and cell types. **C)** Expression of conserved differentially expressed genes in different cell types during bell regeneration in *Aurelia* (left) and *Rhopilema* (right).

**Figure 4.**
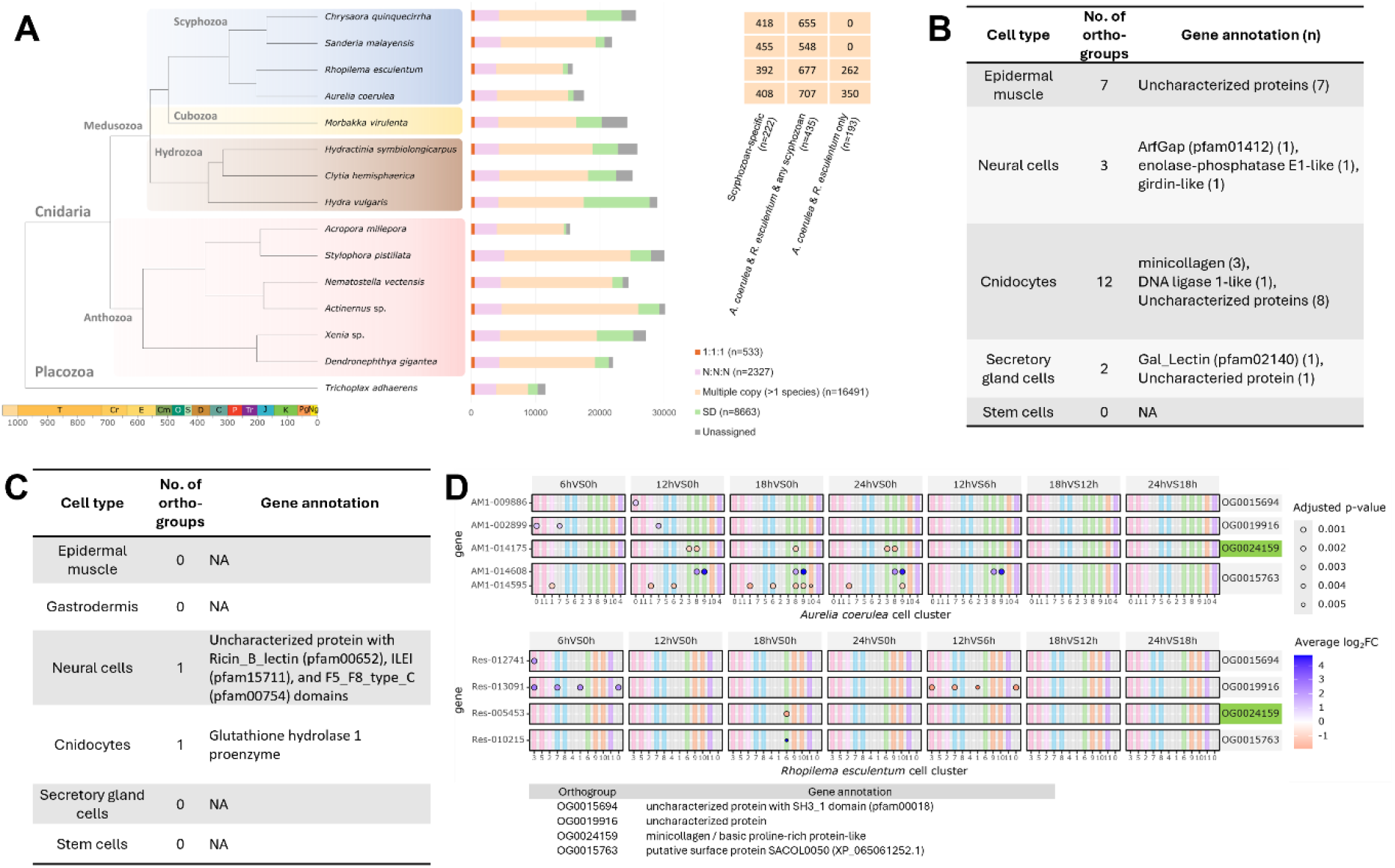
**A)** Summary of genes gained in each investigated metazoan lineage. **B-C)** Scyphozoan-specific orthogroups (B) and toxins (C) with conserved cell-type expression in *Aurelia* and *Rhopilema*. **D)** Dot plot showing scyphozoan lineage-specific genes during bell regeneration in *Aurelia* (upper) and *Rhopilema* (lower). OG0024159 with conserved cnidocyte cell expression is coloured in green.

To understand the genetic regulatory mechanisms underlying the regeneration process in *Aurelia* and *Rhopilema*, gene enrichment analyses were also performed. These revealed that three gene pathways - regulation of actin cytoskeleton, focal adhesion, and leukocyte transendothelial migration, were all upregulated in epidermal muscle in both species at 6h post-amputation (Figure 3B; Supplementary Figure 16). Moreover, temporal differences were also observed for enriched metabolic gene pathways in the neural and secretory gland cells during regeneration in *Aurelia* (6-12h) and *Rhopilema* (18-24h).

To further elucidate the genes involved in jellyfish regeneration, we investigated differentially expressed genes that are conserved between the regeneration process in *Aurelia* and *Rhopilema* (Figure 3C, Supplementary Figure 17). We revealed that transcription factors (*Otx-D, Otx-B*), *TFAP2A*, Wnt/β-catenin signalling pathway genes (*Wnt11, Fzd10, Fzd-like, CK1α*), growth factor signalling pathway genes (*EGF, FGF, FGFR*), and extracellular matrix genes (collagen α-1(I) chain-like, laminin, matrix metalloproteinase-2-like enzymes, dystroglycan adhesion complexes, cytoskeletal regulators filamin-A-like, myosin-10, tropomyosin-2), are all involved in the regeneration process in both *Aurelia* and *Rhopilema*.

### Scyphozoan-specific genes expression during the regeneration process

To understand patterns in gene gains in scyphozoans, we compared orthologous genes across the genome assemblies of four scyphozoans, ten additional cnidarians, and a placozoan outgroup (Supplementary Table 13). Among the conserved scyphozoan-specific orthogroups, 27 orthogroups were expressed in our scRNA-seq data and also showed conserved cell-type expression in both *Aurelia* and *Rhopilema* (Figure 4A-B, Supplementary Data 13). These scyphozoan-specific genes were verified using reciprocal blast searches, amino acid sequence alignments, and phylogenetic tree construction, to confirm they are only present in scyphozoan genomes and not in other investigated metazoan genomes. For 24 of these 27 scyphozoan-specific genes, half were expressed specifically in the cnidocytes (n=12), while the others are expressed in epidermal muscle (n=7), neural cells (n=3), and secretory gland cells (n=2)(Figure 4B, Supplementary information 18-23). For the 12 scyphozoan-specific genes expressed specifically in the cnidocytes, three are minicollagens, including OG0020136 which was previously described (Gold et al 2019). Two scyphozoan-specific genes that were also found in the proteomes were expressed specifically in cnidocytes (n=1) and neural cells (n=1) in *Aurelia* (Figure 4C), but they were expressed in multiple cell types in *Rhopilema* (Supplementary Figure 24-26). Additionally, certain scyphozoan-specific genes are differentially expressed during regeneration in *Aurelia* and *Rhopilema* (Figure 4D; Supplementary Figure 27-28). These include OG0015694 and OG0019916, which both contain the Src homology 3 (SH3_1, Pfam00018) domain and are upregulated in epidermal muscle; minicollagen OG0024159, which is downregulated in cnidocytes; and putative surface protein OG0015763, which is upregulated in cnidocytes.

## Discussion

Here, comparative analyses considering the results of analyses of two newly generated single-cell transcriptome libraries, alongside previously reported findings for other cnidarians, reveal several novel insights into the cell types, conserved gene regulation, and lineage-specific genes involved in cnidarian regeneration processes.

Genomes provide a crucial foundation for developing improved understanding of the conserved and divergent genomic characters present in Cnidaria and Bilateria (Gold et al 2019; Khalturin et al 2019; Nong et al 2020). To gain a better understanding of the genomic sequences contributing at the individual cell level, scRNA-seq has proven an appropriate and powerful tool to study the hallmarks of metazoan evolution (e.g. Sebe-Pedros et al 2018; Shao et al 2020; Chari et al 2021; Ghaddar et al 2023; Cole et al 2024; Dong et al 2024; Mah and Dunn 2024; Baranyk et al 2025). We first show that genome quality and aspects related to genome quality, as well as the gene model prediction algorithms adopted contribute to considerable variation in the mapping rate of scRNA-seq reads in non-model organisms. Hence, we provide a new genome assembly for a male moon jellyfish *Aurelia* from Hong Kong, and test various gene prediction models to establish a more accurate foundation for performing subsequent analyses.

Regeneration in metazoans has been examined anatomically in diverse lineages for centuries (Bideau et al 2021). However, among cnidarians, regeneration in scyphozoans is relatively understudied compared to other lineages, such as anthozoans (e.g. *Exaiptasia* and *Nematostella*), and hydrozoans (e.g. *Clytia* and *Hydra*). By comparing the results of scRNA-seq on regeneration in scyphozoans to those for other cnidarians, we show that regeneration in scyphozoans involves similar cell types and marker genes to other cnidarians. For instance, similar to other cnidarians, myosin chain genes, superviellain (*SVIL*), tropomyosin genes, and calcium signalling regulators are expressed in scyphozoan epidermal muscle cells (Sebé-Pedrós et al, 2018; Nizhnichenko et al, 2022; Tanaka et al. 2018). While the neuronal cell marker gene *ELAV*, advillin, synaptotagmin, tubulin chain, neuropeptide, and voltage-gated ion channels *VGSC* are also expressed in neural cells (Figure 2C, Chuang et al 2018; Geffeney et al 2019; Nakanishi et al 2012; Pascale et al 2008; Takahashi & Takeda, 2015). In scyphozoan gastrodermis cells, *Apolipophorin* expression were identified (Sebé-Pedrós et al 2018), while MUC2 and CREB3L1, which have roles in mucus secretion, were also expressed in the assigned secretory gland cells (Steger et al 2022, Mayorova et al 2019). The gland cell marker gene zinc metalloendopeptidase *ADAMTS* (Sebé-Pedrós et al., 2018) was also found to be expressed in the assigned gland cells of both scyphozoan jellyfishes. For cnidocytes/nematocysts, minicollagens and nematogalectin have been proposed as stinging cell marker genes, and both genes are also preferentially expressed in scyphozoans, in common with other cnidarians (David et al., 2008; Hu et al., 2020; Hwang et al., 2010; Sebé-Pedrós et al., 2018).

Hormone biosynthesis is a crucial component in animal development and evolution, including metamorphosis. In a previous study, sesquiterpenoid hormone production, previously thought to be typical of arthropods only, was discovered in jellyfish and other cnidarians (Nong et al 2020). Nevertheless, knowledge of sesquiterpenoid hormones in cnidarians remains inadequate to date. In the *Aurelia* and *Rhopilema* cell type atlas generated here, sesquiterpenoid biosynthetic pathway genes were found to be expressed in various cell types. In particular, *AACT* (Acetyl-CoA acetyltransferase) in the mevalonate pathway and *ICMT* (Isoprenylcysteine carboxul methyltransferase) downstream to mevalonate pathway, showed strong expression levels. Given none of them were differentially expressed throughout the regeneration in both jellyfish, we suggest that sesquiterpenoid hormones in jellyfish may be involved in life stage transformation and/or reproduction (as in other invertebrates), rather than in playing a key role in scyphozoan regeneration.

Transcription factors are key components of genome function, that have provided fruitful clues into understanding of metazoan regeneration. We identified a suite of conserved transcription factors differentially regulated in both jellyfish considered here, which strongly suggest a conserved genetic regulatory network for regeneration in scyphozoans. Together we identify similar cell type reprogramming during regeneration between the two species considered here. Collectively, these findings suggest that the scyphozoan last common ancestor already possessed established cellular and molecular mechanisms contributing to regeneration.

Lineage-specific genes provide genetic novelty and are frequently identified in metazoan genomes via comparative genomic analysis. Most of our understanding of lineage-specific genes relates to their evolutionary origins, via processes such as gene duplication or horizontal gene transfer (e.g. So et al 2022). The extent to which novel genes contribute to lineage-specific biological processes such as regeneration remains an exciting and largely unexplored area. Here, we first showed that scyphozoan-specific genes are expressed with a conserved pattern in certain cell types, with about half specific to cnidocytes. This explosive cell type is well known to have a distinctive structure and be specific to the Cnidaria for purposes of delivering a sting to other organisms. In addition, we also identified scyphozoan-specific genes that are differentially expressed during regeneration in the two jellyfish species considered here. Overall, these findings imply that certain lineage-specific genes, are under selective pressure to contribute distinct novel morphological structures and biological processes that characterize the diverse lineages that have evolved in the metazoan tree of life.

## Material and Methods

### Animal collection and husbandry

Individuals of *Aurelia* and *Rhopilema* were purchased from a local supplier in Hong Kong. Medusae of both species were maintained and cultured in 30 ppt circulating seawater at 25°C at The Chinese University of Hong Kong. Animals were fed three times weekly with newly hatched brine shrimp *Artemia*.

### DNA extraction of male *Aurelia* and sequencing

The bell and gonad tissues of a male *Aurelia* sp. individual were dissected and snap-frozen in liquid nitrogen. The gonad tissue was used for sex identification via histological examinations with modifications that the tissue was immersed in 4% formalin for 24-hour fixation (García-Rodríguez et al., 2018). The hematoxylin-eosin stained sections were examined under a light microscope (Leica DM500), and the bell tissue was sent to Dovetail Genomics (USA) for high molecular weight DNA isolation. The fragment size and quantity of the isolated HMW DNA was inspected in a 5200 Fragment Analyzer System (Agilent Technologies). A PacBio SMRTbell library was prepared and sequenced on the PacBio Sequel IIe sequencing platform with two SMRT cells to obtain ∼33 Gb data. The Omni-C library and sequencing data was generated by Dovetail Genomics LLC. Briefly, the chromatin was fixed with formaldehyde and then extracted from the nucleus, followed by digestion with DNAse I. Chromatin end repair was performed and a biotinylated bridge adapter was used for proximity ligation of adapter containing ends. Subsequently, crosslink reversal was carried out, followed by DNA purification and removal of biotins that were not ligated to DNA fragments. The library was generated with the use of NEBNext Ultra enzymes and Illumina-compatible adapters. The biotin-containing fragments were retained using streptavidin beads, followed by a PCR enrichment step. The final library was sequenced on an Illumina HiSeqX platform to produce Omni-C data of ∼48X sequencing coverage.

### Genome assembly

*De novo* genome assembly was performed using Hifiasm (version 0.19.8). Haplotypic duplications were identified and removed using purge_dups (version 1.2.5). The primary genome assembly was checked for contamination using BlobTools (version 1.1.1). Proximity ligation data from the Omni-C library was used to scaffold the genome assembly using YaHS (version 1.2a.2). The final assembled genome was soft-masked using redmask (version 0.0.2) and gene models were predicted using funannotate (version 1.8.15) with the parameters “--protein_evidence uniprot_sprot.fasta --cpus 100 --genemark_mode ET --optimize_augustus -- busco_db metazoa --organism other --max_intronlen 350000” as previously described (Nong et al. 2020; So et al 2022).

### Transcriptome sequencing and gene model prediction

Male *Aurelia* gonad was pre-treated with a CTAB solution (1M Tris-HCl, 0.2M EDTA, 5M NaCl, CTAB, 1% PVP and 1% beta-mercaptoethanol) prior to extraction of RNA using TRIzol solution (Invitrogen). Quality control was performed using a Nanodrop spectrophotometer (Thermo Scientific), gel electrophoresis, and an Agilent 2100 Bioanalyzer (Agilent RNA 6000 Nano Kit). Transcriptome sequencing was carried out on Illumina platform (Supplementary Table 1), and data were first processed using trimmomatic (version 0.39) and Kraken (version 2.0.8 with database k2_standard_20210517) and assembled as previously described (Nong et al., 2020; So et al 2022). Other transcriptomic data of *Aurelia* were also downloaded from NCBI database, and mapped to the soft-masked genome using HISAT2 (version 2.2.1). The data generated in this study, as well as sample SRR8040389 have sequence alignment rates of 84.38% and 84.56%, respectively, and were further used for genome annotation using funannotate (version 1.8.15) and BRAKER (v3.0.8).

### Orthologous genes assignment

The longest transcript of gene sets from 15 taxa including 4 scyphozoans (*Aurelia* sp., *Rhopilema esculentum, Chrysaora quinquecirrha* and *Sanderia malayensis*), 1 cubozoan (*Morbakka virulenta*), 3 hydrozoans (*Clytia hemisphaerica, Hydractinia symbiolongicarpus*, and *Hydra vulgaris*), 6 anthozoans (*Actinernus* sp., *Nematostella vectensis, Stylophora pistillata, Acropora millepora, Xenia* sp. and *Dendronephthya gigantea*) and an outgroup from placozoan (*Trichoplax adhaerens*) were used to infer the gene orthology using OrthoFinder v2.5.5 (Emms & Kelly, 2019). *DupGen_finder* (Qiao et al. 2019) and *MCScanX* (Wang et al. 2012) were used to identify syntenic gene pairs between the four analysed scyphozoan genomes. Potential scyphozoan-specific orthogroups were searched for conserved domains in NCBI Conserved Domain Database (Yang et al., 2020; Wang et al., 2023), proteomes of investigated taxa using HMMER (version 3.3.1; cut-off E-value < 10^−5^) (Eddy, 2011), and NCBI nr database. Amino acid sequences were aligned using MAFFT v7.455 (Katoh & Standley, 2013), followed by gene tree constuction with FastTree (Price et al., 2010). Sequence alignment, gene tree and gene synteny were visualized in Jalview v2.11.4.1 (Waterhouse et al., 2009), evolview v3 (Subramanian et al., 2019), and *R* package genomes (Hackl et al., 2021), respectively.

### Jellyfish single cell dissociation and transcriptome sequencing

Bells of individual jellyfish were excised with a stainless-steel trapezium mould (80mm top, 150 mm bottom, 70 mm height). 2mm of tissue was cut along the amputated region at 0-hour, 6-hour, 12-hour, 18-hour and 24-hour post-amputations. Excised tissues were put in trypsin dissociation solution (TrypLE Expression solution, ThermoFisher), and incubated for 30 minutes at 37 °C and 20 minutes at 25 °C for *Rhopilema* and *Aurelia*, respectively. Cells were collected by centrifugation at 600 x g for 5 minutes and resuspended in phosphate-buffered saline (PBS) solution containing 0.04% bovine serum albumin (BSA). Aggregated cells were filtered with Flowmi cell strainer (Bel-Art H-B Instrument). Cells were stained with trypan blue and were counted using hemocytometer to estimate their viability and concentrations. Only cell suspension with cell viability >90% would be further proceeded to barcoding and library construction using the Chromium Next GEM technology, and quantified with High Sensitivity D5000 and D1000 DNA ScreenTape assays (Agilent). Sequencing was carried out using Illumina NovaSeq6000 platform (PE150) with a median depth of 100,000 read pairs per cell.

### Single-cell RNA (scRNA) transcriptome data clustering and cell type atlas identification

ScRNA sequencing data were mapped to the respective reference genomes of *Aurelia* and *Rhopilema* with 10X Genomics Cell Ranger (v7.1.0). Cell-UMI count tables were uploaded to Seurat (v4.4.0), with dead cells, low-quality cells, empty droplets and cell doublets were filtered (Hao et al., 2021). For identification of common cell types, data from all five-timepoints of each jellyfish species were merged and integrated using FindIntegrationAnchors and IntegrateData. ElbowPlot was used to identify the optimal dimensionality, while Clustree_0.4.4 was applied to detect the optimal resolution of clustering (Zappia & Oshlack, 2018). The clustering results were visualized with uniform manifold approximation and projection (UMAP). Published cnidarian single-cell transcriptome data, including *Hydra vulgaris, Clytia hemisphaerica, Xenia* sp., *Stylophora pistillata*, and *Nematostella vectensis* (Chari et al., 2021; Hu et al., 2020; Levy et al., 2021; Sebé-Pedrós et al., 2018; Siebert et al., 2019) were used to define cell types using OrthoFinder (version 2.5.4) with the parameters “- t 138 -a 138 -M msa -S diamond”. Kullback–Leibler divergence (KLD) was calculated for the cell types of *Aurelia, Rhopilema*, and other cnidarians (Levy et al., 2021). Conserved cell markers for different cell populations were identified using Seurat anchors FindAllMarkers (logfc.threshold = 0.25). Representative genes for each cluster were further manually annotated and checked. Marker genes of each cluster were also tested for KEGG, GO, and JOG enrichment analyses (Wu et al., 2021; So et al 2022).

### Pseudotime and differential gene expression analyses during regeneration

Cell differentiation trajectories during the regeneration processes was inferred using R package Monocle3 (Cao et al. 2019), with the germline stem cells of 0 hr of each species being used as root cells for pseudotime analysis. For identification of differentially expressed genes (DEGs) in each cell type, data from 6-hr, 12-hr, 18-hr and 24-hr post-amputation were compared to both 0-hr as well as preceding timepoint of respective species. Functional term annotation for DEGs was performed using EggNOG-mapper v2.1.2 (Cantalapiedra et al. 2021), while enrichment analyses for KEGG pathways and Gene Ontology (GO) were carried out using “clusterProfiler” (Wu et al., 2021). Gene expression changes were visualized with either “ggplot2” package in R or Seurat anchor FeaturePlot or VlnPlot (Villanueva & Chen, 2019).

### Liquid Chromatography-Tandem Mass Spectrometry (LC-MS/MS) on nematocysts

Nematocysts were enriched as previously described (Bloom et al 1998) with the following modifications. Jellyfish samples were first placed in 1:10 (v:v) 35 g/L NaCl at 4 °C, allowing the tissues to autolysis for 4 days. The autolyzed mixture was sequentially filtered through 200 µm and 85 µm cell strainer to remove debris.. Proteins were extracted from enriched nematocysts by dissolving in lysis buffer (6 M urea, 2 M thiourea, 1 mM dithiothreitol (DTT) in 250 mM Tris (pH 7.6) supplemented with Pierce™ protease inhibitor (Thermo Fisher Scientific). Protein samples were alkylated with 5 mM of iodoacetamide for 30 minutes in dark, and further digested overnight at 37 °C with sequencing-grade trypsin (Promega) at a 1:20 ratio. The resulting peptides were desalted with Pierce™ C18 spin columns (Thermo Fisher Scientific) per the manufacturer’s guidelines. LC-MS/MS analysis was performed using a Dionex UltiMate 3000 RSLC nano system interfaced with an Orbitrap Fusion Lumos Tribrid mass spectrometer (Thermo Fisher Scientific). MS and MS/MS scans were acquired in the Orbitrap with a mass resolution of 60,000 and 15,000, respectively. The higher-energy collisional dissociation (HCD) mode was used as the fragmentation mode with 30% collision energy, while the precursor isolation windows were set to 1.6 m/z. Acquired mass spectra were analyzed by Proteome Discoverer version 2.4 with SEQUEST as a search engine. Data were searched against the translated protein sequences of *Aurelia* and *Rhopilema* transcriptomes. Putative toxins were first identified by searching protein sequences against the UniProt animal toxin and venom database using BLASTp (e-value of <1.0 × 10−5) (Jungo et al 2012), with ToxPred2 being used to further exclude proteins with non-toxic physiological functions (Sharma et al 2022).

## Data and code availability

The raw reads generated in this study have been deposited in the NCBI database under BioProject accession numbers PRJNA1229829 (PacBio HiFi and Omni-C reads of *Aurelia*), PRJNA1049062 (single cell sequencing data of *Aurelia*) and PRJNA1045829 (single cell data of *Rhopilema*). The genome and genome annotation files have been deposited in the Figshare data set (https://figshare.com/s/f89fdf550cf68d97a541).

## Acknowledgements

This work was supported by the Hong Kong Research Grant Council General Research Fund (14103823, 14100919) and Collaborative Research Fund (C4015-20EF), and the TUYF Charitable Trust.

## Author contributions

YL, THL, and JT carried out the single-cell isolation. YL, STS, WN, YX carried out the single-cell analyses. WN carried out the gene annotation. WLS, TCNL, SMN carried out the toxin proteomic analyses. OJ and JZ assembled the genome. HYY ensured the sequencing project management and animal husbandry. APYC, KFL, AJ, ZPK, WGB, AH, YW, TFC, and SMN contributed the discussion of project at different stages. JHLH designed and coordinated the project. YL, STSL, WN, and JHLH wrote the initial manuscript; all authors revised and contributed to the final version of the text.

## References

Abrams, M. J., Basinger, T., Yuan, W., Guo, C.-L., & Goentoro, L. (2015). Self-repairing symmetry in jellyfish through mechanically driven reorganization. Proceedings of the National Academy of Sciences, 112(26), E3365–E3373.

Abrams, M. J., Tan, F. H., Li, Y., Basinger, T., Heithe, M. L., Sarma, A., Lee, I. T., Condiotte, Z. J., Raffiee, M., Dabiri, J. O., Gold, D. A., & Goentoro, L. (2021). A conserved strategy for inducing appendage regeneration in moon jellyfish, Drosophila, and mice. eLife, 10, e65092. 10.7554/eLife.65092

Baranyk, J., Malir, K., Silva, M. A., Rieck, S., Scheve, G., & Nakanishi, N. (2025). Structural, molecular and developmental evidence for cell-type diversity in cnidarian mechanosensory neurons. Nature Communications, 16(1), 1514.

Baumgarten, S., Simakov, O., Esherick, L. Y., Liew, Y. J., Lehnert, E. M., Michell, C. T., Li, Y., Hambleton, E. A., Guse, A., Oates, M. E., & others. (2015). The genome of Aiptasia, a sea anemone model for coral symbiosis. Proceedings of the National Academy of Sciences, 112(38), 11893–11898.

Bideau, L., Kerner, P., Hui, J., Vervoort, M., & Gazave, E. (2021). Animal regeneration in the era of transcriptomics. Cellular and Molecular Life Sciences, 78(8), 3941–3956.

Cantalapiedra, C. P., Herņandez-Plaza, A., Letunic, I., Bork, P., & Huerta-Cepas, J. (2021). eggNOG-mapper v2: Functional Annotation, Orthology Assignments, and Domain Prediction at the Metagenomic Scale. Molecular Biology and Evolution, 38(12), 5825–5829. 10.1093/MOLBEV/MSAB293

Cao, J., Spielmann, M., Qiu, X., Huang, X., Ibrahim, D. M., Hill, A. J., Zhang, F., Mundlos, S., Christiansen, L., Steemers, F. J., & others. (2019). The single-cell transcriptional landscape of mammalian organogenesis. Nature, 566(7745), 496–502.

Chang, E. S., Neuhof, M., Rubinstein, N. D., Diamant, A., Philippe, H., Huchon, D., & Cartwright, P. (2015). Genomic insights into the evolutionary origin of Myxozoa within Cnidaria. Proceedings of the National Academy of Sciences, 112(48), 14912–14917.

Chapman, J. A., Kirkness, E. F., Simakov, O., Hampson, S. E., Mitros, T., Weinmaier, T., Rattei, T., Balasubramanian, P. G., Borman, J., Busam, D., & others. (2010). The dynamic genome of Hydra. Nature, 464(7288), 592–596.

Chari, T., Weissbourd, B., Gehring, J., Ferraioli, A., Leclère, L., Herl, M., Gao, F., Chevalier, S., Copley, R. R., Houliston, E., Anderson, D. J., & Pachter, L. (2021). Whole-animal multiplexed single-cell RNA-seq reveals transcriptional shifts across Clytia medusa cell types. Science Advances, 7(48), 1683. 10.1126/SCIADV.ABH1683

Chuang, Y.-C., Lee, C.-H., Sun, W.-H., & Chen, C.-C. (2018). Involvement of advillin in somatosensory neuron subtype-specific axon regeneration and neuropathic pain. Proceedings of the National Academy of Sciences, 115(36), E8557–E8566.

Cole, A. G., Steger, J., Hagauer, J., Denner, A., Ferrer Murguia, P., Knabl, P., Narayanaswamy, S., Wick, B., Montenegro, J. D., & Technau, U. (2024). Updated single cell reference atlas for the starlet anemone Nematostella vectensis. Frontiers in Zoology, 21(1), 8.

Curtis, S. K., & Cowden, R. R. (1974). Some aspects of regeneration in the scyphistoma of Cassiopea (Class Scyphozoa) as revealed by the use of antimetabolites and microspectrophotometry. American Zoologist, 14(2), 851–866.

David, C. N., Özbek, S., Adamczyk, P., Meier, S., Pauly, B., Chapman, J., Hwang, J. S., Gojobori, T., & Holstein, T. W. (2008). Evolution of complex structures: Minicollagens shape the cnidarian nematocyst. Trends in Genetics, 24(9), 431–438. 10.1016/J.TIG.2008.07.001

Dong, Z., Wang, F., Liu, Y., Li, Y., Yu, H., Peng, S., Sun, T., Qu, M., Sun, K., Wang, L., & others. (2024). Genomic and single-cell analyses reveal genetic signatures of swimming pattern and diapause strategy in jellyfish. Nature Communications, 15(1), 936.

DuBuc, T. Q., Traylor-Knowles, N., & Martindale, M. Q. (2014). Initiating a regenerative response; cellular and molecular features of wound healing in the cnidarian Nematostella vectensis. BMC Biology, 12, 1–21.

Eddy, S. R. (2011). Accelerated profile HMM searches. PLoS Computational Biology, 7(10), e1002195.

Emms, D. M., & Kelly, S. (2019). OrthoFinder: Phylogenetic orthology inference for comparative genomics. Genome Biology, 20, 1–14.

Fox, R. M., Hanlon, C. D., & Andrew, D. J. (2010). The CrebA/Creb3-like transcription factors are major and direct regulators of secretory capacity. Journal of Cell Biology, 191(3), 479–492.

Gamero-Mora, E., Halbauer, R., Bartsch, V., Stampar, S. N., & Morandini, A. C. (2019). Regenerative capacity of the upside-down jellyfish Cassiopea xamachana. Zoological Studies, 58, e37.

García-Rodríguez, J., Lewis Ames, C., Marian, J. E. A., & Marques, A. C. (2018). Gonadal histology of box jellyfish (Cnidaria: Cubozoa) reveals variation between internal fertilizing species Alatina alata (Alatinidae) and Copula sivickisi (Tripedaliidae). Journal of Morphology, 279(6), 841–856.

Geffeney, S. L., Williams, B. L., Rosenthal, J. J., Birk, M. A., Felkins, J., Wisell, C. M., Curry, E. R., & Hanifin, C. T. (2019). Convergent and parallel evolution in a voltage-gated sodium channel underlies TTX-resistance in the greater blue-ringed octopus: Hapalochlaena lunulata. Toxicon, 170, 77–84.

Ghaddar, B., Blaser, M. J., & De, S. (2023). Denoising sparse microbial signals from single-cell sequencing of mammalian host tissues. Nature Computational Science, 3(9), 741–747.

Gold, D. A., Katsuki, T., Li, Y., Yan, X., Regulski, M., Ibberson, D., Holstein, T., Steele, R. E., Jacobs, D. K., & Greenspan, R. J. (2019). The genome of the jellyfish Aurelia and the evolution of animal complexity. Nature Ecology & Evolution, 3(1), 96–104.

Gold, D. A., Lau, C. L. F., Fuong, H., Kao, G., Hartenstein, V., & Jacobs, D. K. (2019). Mechanisms of cnidocyte development in the moon jellyfish Aurelia. Evolution & Development, 21(2), 72–81.

Gurtner, G. C., Werner, S., Barrandon, Y., & Longaker, M. T. (2008). Wound repair and regeneration. Nature, 453(7193), 314–321.

Hackl, T., Duponchel, S., Barenhoff, K., Weinmann, A., & Fischer, M. G. (2021). Virophages and retrotransposons colonize the genomes of a heterotrophic flagellate. Elife, 10, e72674.

Hao, Y., Hao, S., Andersen-Nissen, E., Mauck, W. M., Zheng, S., Butler, A., Lee, M. J., Wilk, A. J., Darby, C., Zager, M., & others. (2021). Integrated analysis of multimodal single-cell data. Cell, 184(13), 3573–3587.

Hu, M., Zheng, X., Fan, C. M., & Zheng, Y. (2020). Lineage dynamics of the endosymbiotic cell type in the soft coral Xenia. Nature, 582(7813), 534–538. 10.1038/s41586-020-2385-7

Hwang, J. S., Takaku, Y., Momose, T., Adamczyk, P., Özbek, S., Ikeo, K., Khalturin, K., Hemmrich, G., Bosch, T. C. G., Holstein, T. W., David, C. N., & Gojobori, T. (2010). Nematogalectin, a nematocyst protein with GlyXY and galectin domains, demonstrates nematocyte-specific alternative splicing in Hydra. Proceedings of the National Academy of Sciences, 107(43), 18539–18544. 10.1073/pnas.1003256107

Jungo, F., Bougueleret, L., Xenarios, I., & Poux, S. (2012). The UniProtKB/Swiss-Prot Tox-Prot program: A central hub of integrated venom protein data. Toxicon, 60(4), 551–557.

Katoh, K., & Standley, D. M. (2013). MAFFT multiple sequence alignment software version 7: Improvements in performance and usability. Molecular Biology and Evolution, 30(4), 772–780.

Khalturin, K., Shinzato, C., Khalturina, M., Hamada, M., Fujie, M., Koyanagi, R., Kanda, M., Goto, H., Anton-Erxleben, F., Toyokawa, M., & others. (2019). Medusozoan genomes inform the evolution of the jellyfish body plan. Nature Ecology & Evolution, 3(5), 811–822.

Layden, M. J., Rentzsch, F., & Röttinger, E. (2016). The rise of the starlet sea anemone Nematostella vectensis as a model system to investigate development and regeneration. Wiley Interdisciplinary Reviews: Developmental Biology, 5(4), 408–428.

Leclère, L., Horin, C., Chevalier, S., Lapébie, P., Dru, P., Peron, S., Jager, M., Condamine, T., Pottin, K., Romano, S., & others. (2019). The genome of the jellyfish Clytia hemisphaerica and the evolution of the cnidarian life-cycle. Nature Ecology & Evolution, 3(5), 801–810.

Levy, S., Elek, A., Grau-Bové, X., Menéndez-Bravo, S., Iglesias, M., Tanay, A., Mass, T., & Sebé-Pedrós, A. (2021). A stony coral cell atlas illuminates the molecular and cellular basis of coral symbiosis, calcification, and immunity. Cell, 184(11), 2973–2987.

Liu, Y., Zhou, Q., Wang, Y., Luo, L., Yang, J., Yang, L., Liu, M., Li, Y., Qian, T., Zheng, Y., & others. (2015). Gekko japonicus genome reveals evolution of adhesive toe pads and tail regeneration. Nature Communications, 6(1), 10033.

Mah, J. L., & Dunn, C. W. (2024). Cell type evolution reconstruction across species through cell phylogenies of single-cell RNA sequencing data. Nature Ecology & Evolution, 8(2), 325–338.

Mayorova, T. D., Hammar, K., Winters, C. A., Reese, T. S., & Smith, C. L. (2019). The ventral epithelium of Trichoplax adhaerens deploys in distinct patterns cells that secrete digestive enzymes, mucus or diverse neuropeptides. Biology Open, 8(8), bio045674.

Nakanishi, N., Renfer, E., Technau, U., & Rentzsch, F. (2012). Nervous systems of the sea anemone Nematostella vectensis are generated by ectoderm and endoderm and shaped by distinct mechanisms. Development, 139(2), 347–357. 10.1242/DEV.071902

Nizhnichenko, V. A., Boyko, A. V., Ginanova, T. T., & Dolmatov, I. Y. (2022). Muscle Regeneration in Holothurians without the Upregulation of Muscle Genes. International Journal of Molecular Sciences, 23(24), Article 24. 10.3390/ijms232416037

Nong, W., Cao, J., Li, Y., Qu, Z., Sun, J., Swale, T., Yip, H. Y., Qian, P. Y., Qiu, J.-W., Kwan, H. S., & others. (2020). Jellyfish genomes reveal distinct homeobox gene clusters and conservation of small RNA processing. Nature Communications, 11(1), 3051.

Nowoshilow, S., Schloissnig, S., Fei, J.-F., Dahl, A., Pang, A. W., Pippel, M., Winkler, S., Hastie, A. R., Young, G., Roscito, J. G., & others. (2018). The axolotl genome and the evolution of key tissue formation regulators. Nature, 554(7690), 50–55.

Price, M. N., Dehal, P. S., & Arkin, A. P. (2010). FastTree 2–approximately maximum-likelihood trees for large alignments. PloS One, 5(3), e9490.

Putnam, N. H., Srivastava, M., Hellsten, U., Dirks, B., Chapman, J., Salamov, A., Terry, A., Shapiro, H., Lindquist, E., Kapitonov, V. V., & others. (2007). Sea anemone genome reveals ancestral eumetazoan gene repertoire and genomic organization. Science, 317(5834), 86–94.

Qiao, X., Li, Q., Yin, H., Qi, K., Li, L., Wang, R., Zhang, S., & Paterson, A. H. (2019). Gene duplication and evolution in recurring polyploidization–diploidization cycles in plants. Genome Biology, 20, 1–23.

Sebé-Pedrós, A., Saudemont, B., Chomsky, E., Plessier, F., Mailhé, M.-P. P., Renno, J., Loe-Mie, Y., Lifshitz, A., Mukamel, Z., Schmutz, S., Novault, S., Steinmetz, P. R. H., Spitz, F., Tanay, A., & Marlow, H. (2018). Cnidarian cell type diversity and regulation revealed by whole-organism single-cell RNA-Seq. Cell, 173(6), 1520–1534. 10.1016/j.cell.2018.05.019

Shao, Y., Wang, X.-B., Zhang, J.-J., Li, M.-L., Wu, S.-S., Ma, X.-Y., Wang, X., Zhao, H.-F., Li, Y., Zhu, H. H., Irwin, D. M., Wang, D.-P., Zhang, G.-J., Ruan, J., & Wu, D.-D. (2020). Genome and single-cell RNA-sequencing of the earthworm Eisenia andrei identifies cellular mechanisms underlying regeneration. Nature Communications, 11(1), Article 1. 10.1038/s41467-020-16454-8

Sharma, N., Naorem, L. D., Jain, S., & Raghava, G. P. (2022). ToxinPred2: An improved method for predicting toxicity of proteins. Briefings in Bioinformatics, 23(5), bbac174.

Shinzato, C., Shoguchi, E., Kawashima, T., Hamada, M., Hisata, K., Tanaka, M., Fujie, M., Fujiwara, M., Koyanagi, R., Ikuta, T., & others. (2011). Using the Acropora digitifera genome to understand coral responses to environmental change. Nature, 476(7360), 320–323.

Shum, C. W., Nong, W., So, W. L., Li, Y., Qu, Z., Yip, H. Y., Swale, T., Ang, P. O., Chan, K. M., Chan, T. F., & others. (2022). Genome of the sea anemone Exaiptasia pallida and transcriptome profiles during tentacle regeneration. Frontiers in Cell and Developmental Biology, 10, 900321.

Siebert, S., Farrell, J. A., Cazet, J. F., Abeykoon, Y., Primack, A. S., Schnitzler, C. E., & Juliano, C. E. (2019). Stem cell differentiation trajectories in Hydra resolved at single-cell resolution. Science, 365(6451). 10.1126/science.aav9314

Sinigaglia, C., Peron, S., Eichelbrenner, J., Chevalier, S., Steger, J., Barreau, C., Houliston, E., & Leclère, L. (2020). Pattern regulation in a regenerating jellyfish. Elife, 9, e54868.

So, W. L., Nong, W., Xie, Y., Baril, T., Ma, H., Qu, Z., Haimovitz, J., Swale, T., Gaitan-Espitia, J. D., Lau, K. F., & others. (2022). Myriapod genomes reveal ancestral horizontal gene transfer and hormonal gene loss in millipedes. Nature Communications, 13(1), 3010.

Steger, J., Cole, A. G., Denner, A., Lebedeva, T., Genikhovich, G., Ries, A., Reischl, R., Taudes, E., Lassnig, M., & Technau, U. (2022). Single-cell transcriptomics identifies conserved regulators of neuroglandular lineages. Cell Reports, 40(12).

Steinberg, S. N. (1963). The regeneration of whole polyps from ectodermal fragments of scyphistoma larvae of Aurelia aurita. The Biological Bulletin, 124(3), 337–343.

Stierwald, M., Yanze, N., Bamert, R. P., Kammermeier, L., & Schmid, V. (2004). The Sine oculis/Six class family of homeobox genes in jellyfish with and without eyes: Development and eye regeneration. Developmental Biology, 274(1), 70–81.

Subramanian, B., Gao, S., Lercher, M. J., Hu, S., & Chen, W.-H. (2019). Evolview v3: A webserver for visualization, annotation, and management of phylogenetic trees. Nucleic Acids Research, 47(W1), W270–W275.

Takahashi, T., & Takeda, N. (2015). Insight into the molecular and functional diversity of cnidarian neuropeptides. International Journal of Molecular Sciences, 16(2), 2610–2625. 10.3390/ijms16022610

Tanaka, H., Ishimaru, S., Nagatsuka, Y., & Ohashi, K. (2018). Smooth muscle-like Ca2+-regulation of actin–myosin interaction in adult jellyfish striated muscle. Scientific Reports, 8(1), 7776. 10.1038/s41598-018-24817-x

Vila-Farré, M., Rozanski, A., Ivanković, M., Cleland, J., Brand, J. N., Thalen, F., Grohme, M. A., von Kannen, S., Grosbusch, A. L., Vu, H. T.-K., & others. (2023). Evolutionary dynamics of whole-body regeneration across planarian flatworms. Nature Ecology & Evolution, 7(12), 2108–2124.

Vogg, M. C., Beccari, L., Iglesias Ollé, L., Rampon, C., Vriz, S., Perruchoud, C., Wenger, Y., & Galliot, B. (2019). An evolutionarily-conserved Wnt3/β-catenin/Sp5 feedback loop restricts head organizer activity in Hydra. Nature Communications, 10(1), 312.

Wang, J., Chitsaz, F., Derbyshire, M. K., Gonzales, N. R., Gwadz, M., Lu, S., Marchler, G. H., Song, J. S., Thanki, N., Yamashita, R. A., & others. (2023). The conserved domain database in 2023. Nucleic Acids Research, 51(D1), D384–D388.

Wang, Y., Tang, H., DeBarry, J. D., Tan, X., Li, J., Wang, X., Lee, T., Jin, H., Marler, B., Guo, H., & others. (2012). MCScanX: a toolkit for detection and evolutionary analysis of gene synteny and collinearity. Nucleic Acids Research, 40(7), e49–e49.

Waterhouse, A. M., Procter, J. B., Martin, D. M., Clamp, M., & Barton, G. J. (2009). Jalview Version 2—A multiple sequence alignment editor and analysis workbench. Bioinformatics, 25(9), 1189–1191.

Wu, T., Hu, E., Xu, S., Chen, M., Guo, P., Dai, Z., Feng, T., Zhou, L., Tang, W., Zhan, L., & others. (2021). clusterProfiler 4.0: A universal enrichment tool for interpreting omics data. The Innovation, 2(3).

Yang, M., Derbyshire, M. K., Yamashita, R. A., & Marchler-Bauer, A. (2020). NCBI’s conserved domain database and tools for protein domain analysis. Current Protocols in Bioinformatics, 69(1), e90.

Yu, Y., Nong, W., So, W. L., Xie, Y., Yip, H. Y., Haimovitz, J., Swale, T., Baker, D. M., Bendena, W. G., Chan, T. F., & others. (2022). Genome of elegance coral Catalaphyllia jardinei (Euphylliidae). Frontiers in Marine Science, 9, 991391.

Zappia, L., & Oshlack, A. (2018). Clustering trees: A visualization for evaluating clusterings at multiple resolutions. Gigascience, 7(7), giy083.

Zeleny, C. (1907). The effect of degree of injury, successive injury and functional activity upon regeneration in the scyphomedusan, Cassiopea xamachana. Journal of Experimental Zoology, 5, 265–274.

